# Encoding of facial features by single neurons in the human amygdala and hippocampus

**DOI:** 10.1101/2020.11.02.363853

**Authors:** Runnan Cao, Xin Li, Nicholas J. Brandmeir, Shuo Wang

## Abstract

The human amygdala and hippocampus play a key role in face processing. However, it has been unknown how the neurons in the human amygdala and hippocampus encode facial feature information and directs eye movements to salient facial features such as the eyes and mouth. In this study, we identified a population of neurons that differentiated fixations on the eyes vs. mouth. The response of these feature-selective neurons was not dependent on fixation order, and eye-preferring and mouth-preferring neurons were not of different neuronal types. We found another population of neurons that differentiated saccades to the eyes vs. mouth. Population decoding confirmed our results and further revealed the temporal dynamics of face feature coding. Interestingly, we found that the amygdala and hippocampus played a different role in encoding face features. Lastly, we revealed two functional roles of feature-selective neurons that they encoded the salient region for face recognition and they encoded perceived social trait judgment. Together, we revealed and characterized a new class of neurons that encoded facial features. These neurons may play an important role in social perception and recognition of faces.

## Introduction

The human amygdala and hippocampus have long been associated with a key role in processing faces (Adolphs 2008, Fried et al 1997, Kreiman et al 2000, Mende-Siedlecki et al 2013, Quian Quiroga et al 2005, Viskontas et al 2009). Single neurons in the human amygdala and hippocampus are not only visually selective to faces (Kreiman et al 2000) and facial emotions (Fried et al 1997), but also encode face identities (Quian Quiroga et al 2005) and personally relevant identities (Viskontas et al 2009). Furthermore, amygdala neurons encode subjective judgment of facial emotions (Wang et al 2014) and perceptual decisions to discern one person from another (Quian Quiroga et al 2014), rather than simply process their facial features. A recent study using a unique combination of single-neuron recordings, functional magnetic resonance imaging (fMRI), and patients with focal amygdala lesions has shown that the human amygdala parametrically encodes the intensity of specific facial emotions and their categorical ambiguity (Wang et al 2017). Consistent with these single-neuron studies, intracranial field potentials in the amygdala recorded from implanted depth electrodes show stronger gamma-band activity to faces than to houses or to scrambled faces (Sato et al 2012) and modulation by emotion and attention (Pourtois et al 2010). Findings from human studies are further complemented by monkey studies: a high-resolution fMRI study in monkeys found greater activation in the amygdala to images of monkey faces and bodies than to their scrambled versions (Logothetis et al 1999). Electrophysiological recordings in monkeys (Leonard et al 1985, Rolls 1984) have found single neurons that respond not only to faces, but also to face identities and facial expressions (Gothard et al 2007, Hoffman et al 2007).

However, the mechanisms by which the human amygdala and hippocampus process facial feature information and directs eye movements to salient facial features remain unclear. Neuroimaging studies have shown that amygdala activation predicts gaze direction (Gamer & Büchel 2009) and single neurons in the human amygdala have been found to encode whole faces compared to piecemeal faces (Rutishauser et al 2011), indicating that amygdala neurons may encode facial parts as well as holistic facial features. Furthermore, neurons in the monkey amygdala have been shown to encode not only the eyes but also the gaze direction when viewing a monkey face as well as eye contact with the viewed monkey (Hoffman et al 2007, Mosher et al 2014).

In this study, we used natural face stimuli to study the neural correlates of human eye movements when participants viewed faces. We identified a subset of neurons in the human MTL that differentiated fixations on different facial features and characterized the response of these neurons in detail. Interestingly, we found that these facial feature selective neurons also encoded the saliency of facial features for face identification and they were correlated with social trait judgment.

## Results

### Behavior

We recorded single neurons using implanted micro depth electrodes in the human amygdala and hippocampus (MTL areas) while neurosurgical patients performed a one-back task using real-world images of famous faces (**Fig. 1A**; accuracy = 75.7±5.28% [mean±SD across sessions]). Five patients undergoing epilepsy monitoring had normal basic ability to discriminate faces and they underwent 16 sessions in total (3 sessions were excluded that had fewer than 10 fixations onto each region of interest (ROI), resulting in a total of 13 sessions for further analysis; **Table S1**). Participants viewed 500 natural face images of 50 celebrities (10 faces per identity; **Fig. 1B**). We found that on average, 22.01%±13.10% of fixations were on the eyes and 13.79% ±12.95% of fixations were on the mouth (**Fig. 1B, C**). Furthermore, we found that participants had 16.50%±9.28% of saccades to the eyes and 13.80%±8.49% of saccades to the mouth (**Fig. 1B, D**). Interestingly, among the first fixations, participants tended to fixate 10% more onto the eyes than the mouth (**Fig. E**; two-tailed paired *t*-test: *t*(11) = 2.39, P = 0.036), suggesting that they sampled the eyes earlier than the mouth.

**Fig. 1.**
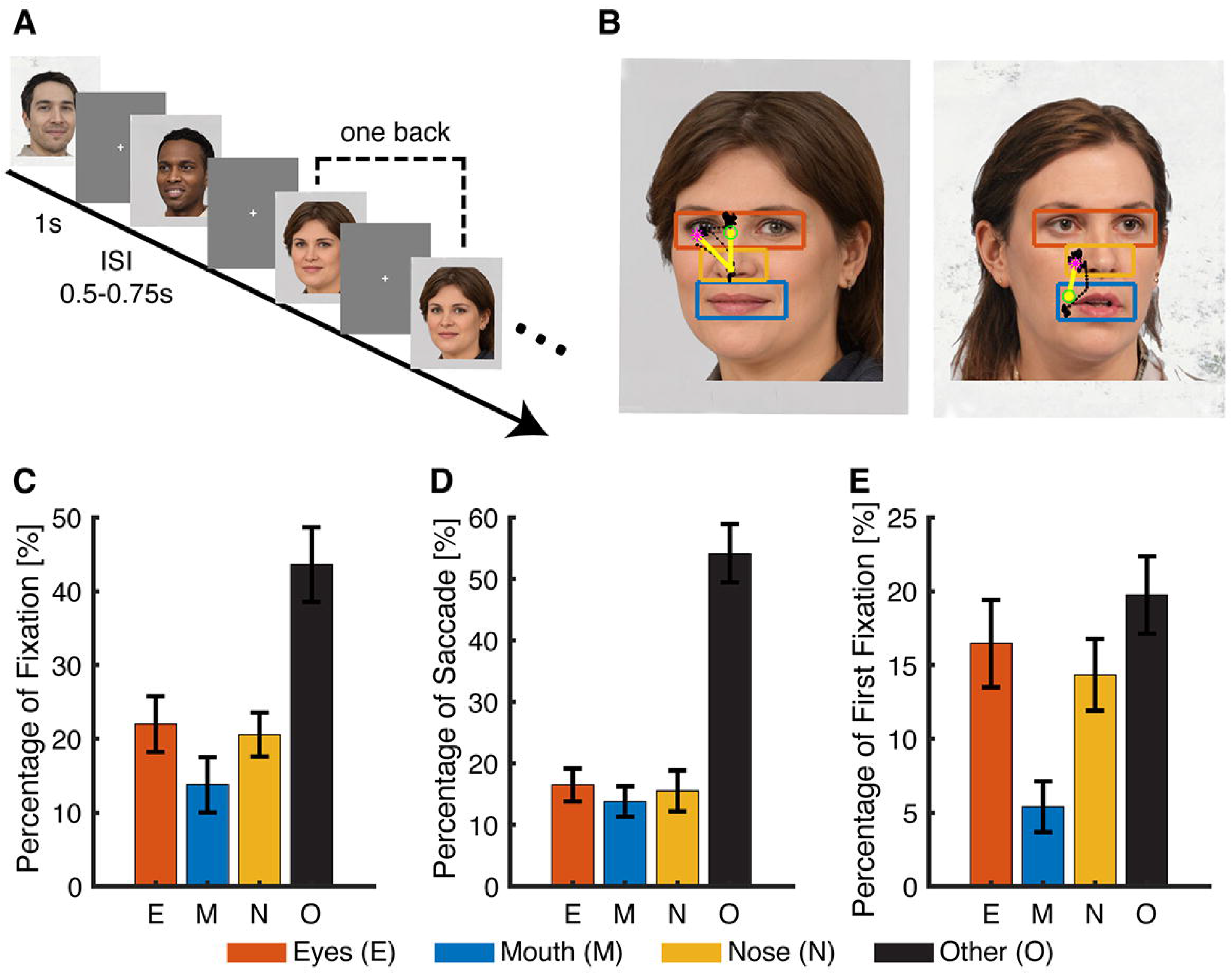
Behavior. **(A)** Task. We employed a one-back task in which patients responded whenever an identical famous face was repeated. Each face was presented for 1s, followed by a jittered inter-stimulus-interval (ISI) of 0.5 to 0.75 seconds. **(B)** Sample stimuli. Regions of interest (ROIs) were detected using computer vision (not shown to participants). Each yellow dot represents a fixation. Green circle: first fixation. Magenta asterisk: last fixation. Yellow line: saccades. Black dot: raw gaze position. **(C)** Percentage of fixations for each ROI. **(D)** Percentage of saccades to each ROI. **(E)** Percentage of first fixation onto each ROI. Error bars denote ±SEM across sessions. E: eyes. M: mouth. N: nose. O: other (all other parts of the image, including hair, neck, background, etc.). Note: To avoid copyright problem, face images displaying in Fig. 1 and Fig. 6 were artificial-intelligence-synthesized from https://generated.photos/faces/natural/ and have no real identities.

### Feature-selective neurons that discriminated fixations on eyes vs. mouth

We isolated 422 single units across all sessions; and of these, 365 units had an overall firing rate greater than 0.15 Hz and we restricted our analysis to this subset of units, which included 178 units from the amygdala, 139 units form the anterior hippocampus, and 48 units from the posterior hippocampus (**Table S1**; see **Fig. S1** for assessment of spike sorting quality).

To investigate the neural encoding of facial features, we first analyzed the response of each neuron between fixations on the eyes and mouth. We aligned neuronal responses at fixation onset and used the mean firing rate in a time window starting 200 ms before fixation onset and ending 200 ms after fixation offset (next saccade onset) to calculate statistics. The duration of this window was on average 814.7±105.8 ms (mean±SD across sessions). We identified 74/365 neurons (20.27%; binomial P < 10^−20^; **Table S1**) that had a response differing significantly between fixations on the eyes vs. the mouth (two-tailed *t*-test, P < 0.05). We identified two types of such feature-selective neurons: one type had a greater response to the eyes relative to the mouth (“eye-preferring”; 55/74 neurons [74%]; see **Fig. 2A, B** for individual examples and **Fig. 2E** for group results) and the second type had a greater response to the mouth relative to the eyes (“mouth-preferring”; 19/74 neurons [26%]; see **Fig. 2C, D** for individual examples and **Fig. 2F** for group results). To investigate the relationship between the response of these feature-selective neurons and their behavior, we quantified the response of feature-selective neurons during individual fixations using a fixation-selectivity index (FSI; see **Eq. 1,2** and **Methods**). As expected, the FSI for feature-selective neurons was significantly larger during fixations on the eyes compared to fixations on the mouth (two-tailed two-sample Kolmogorov-Smirnov [KS] test, KS = 0.26, P < 10^−20^; **Fig. 2G**). This confirms that the single-fixation response of feature-selective neurons is strong enough to allow single-fixation analysis (see **Fig. 2H** for ROC analysis). Permutation tests by shuffling the label of eyes and mouth further confirmed our results: feature-selective neurons (48.15%±5.41%, mean±SD across neurons) had a significantly higher FSI compared to chance (13.90%±1.54%, permutation P < 0.001), whereas the FSI of all non-feature-selective neurons (20.2%±1.49%) was not significantly above chance (19.50% ±1.35%, permutation P = 0.28). Furthermore, we found that feature-selective neurons not only differentiated between the eyes vs. the mouth, but also differentiated the eyes and mouth from other facial parts (i.e., the “nose” and “other” [all other areas of the entire image, including hair, neck, background, etc] ROIs; KS-test: all Ps < 10^−11^; **Fig. 2G**), suggesting that feature-selective neurons were specifically tuned for the eyes and mouth.

**Fig. 2.**
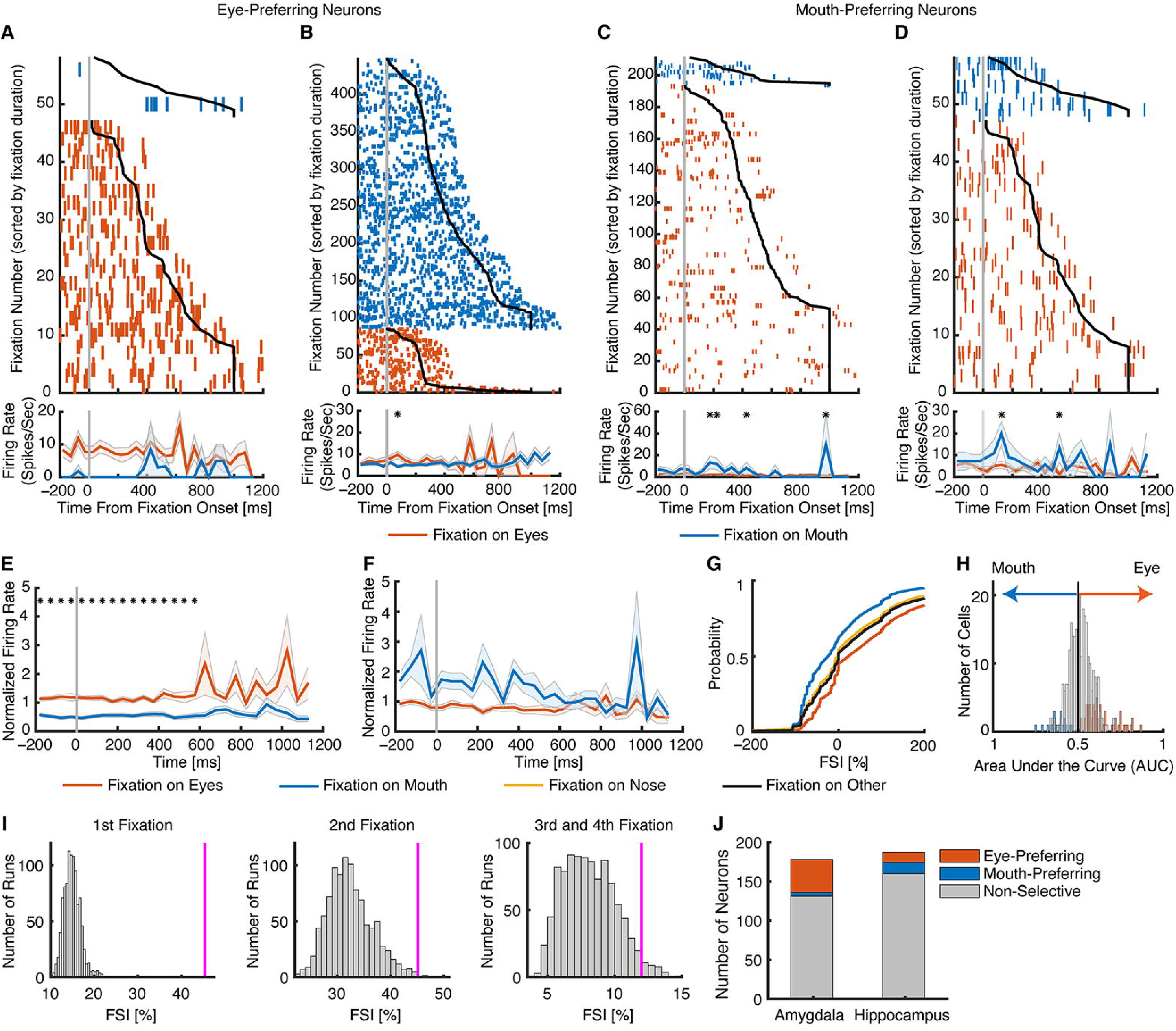
Neurons that differentiate fixations on the eyes vs. the mouth. **(A-D)** Fixation-aligned individual examples. **(A, B)** Neurons that had a greater firing rate when fixating on the eyes compared to the mouth (selection by two-tailed *t*-test in a time window of −200 ms before fixation onset to 200 ms after fixation offset: both Ps < 0.001). **(C, D)** Neurons that had a greater firing rate when fixating on the mouth compared to the eyes (both Ps < 0.01). Fixations are sorted by fixation duration (black line shows start of the next saccade). Fixation onset is t = 0. Asterisk indicates a significant difference between fixations on the eyes and mouth in that bin (P < 0.05, two-tailed *t*-test, after Bonferroni correction; bin size = 50 ms). **(E-J)** Population summary of all feature-selective neurons. **(E)** Average normalized firing rate of eye-preferring neurons (n = 55). **(F)** Average normalized firing rate of mouth-preferring neurons (n = 19). Shaded area denotes ±SEM across neurons. Asterisk indicates a significant difference between the conditions in that bin (P < 0.05, two-tailed *t*-test, after Bonferroni correction). **(G)** Single-fixation analysis using the fixation-selectivity index (FSI; **Methods**). Shown is the cumulative distribution of the single-fixation response of fixation-aligned eye- and mouth-preferring neurons for fixations on the eyes and mouth (n = 74 neurons). **(H)** Population summary using ROC analysis. Shown are histograms of AUC values of eye-preferring neurons (red), mouth-preferring neurons (blue), and neurons that are neither eye-preferring nor mouth-preferring (gray). **(I)** FSI for fixation serial order. The magenta line indicates the observed mean FSI. The null distribution of mean FSI (shown in gray histogram) was calculated by permutation tests of shuffling the labels of fixations on the eyes and mouth (1000 runs). **(J)** The number of neurons in the amygdala and hippocampus from which recordings were made. Stacked bar shows eye-preferring neurons (red), mouth-preferring neurons (blue), and non-feature-selective neurons (gray).

Because participants started viewing the faces from the image center given a preceding central fixation cross, the serial order of fixation onto each ROI might confound feature-selectivity and stimulus-evoked neuronal response (i.e., stimulus-evoked response might result in a greater response for later fixations). In a control analysis of fixation serial order, we found that feature-selective neurons still had a significantly above-chance FSI using the first fixation (45.37% ±47.84%, permutation P < 0.001), second fixation (45.14%±67.73%, permutation P = 0.005), and combined third and forth fixations (12.00%±29.96%, permutation P = 0.033; **Fig. 2I**) onto the eyes vs. the mouth alone, suggesting that our observed feature-selectivity could not be attributed to fixation serial order and stimulus-evoked neuronal response. Lastly, the difference preceding fixation onset (**Fig. 2E, F**) was likely due to saccade planning and multiple fixations in the same ROI. However, we found qualitatively the same results when using different time windows to select feature-selective neurons (e.g., excluding time intervals before and/or after fixations).

Taken together, our results have shown that a subset of MTL neurons encode facial features by discriminating fixations onto the eyes vs. the mouth.

### Comparison of cell types between eye-preferring and mouth-preferring neurons

It is worth noting that for both eye-preferring neurons and mouth-preferring neurons, the difference was primarily driven by fixations on the mouth: eye-preferring neurons had a decreasing firing rate for fixations on the mouth while mouth-preferring neurons had an increasing firing rate for fixations on the mouth, whereas the firing rate for fixations on the eyes remained relatively constant throughout the fixation period for both eye-preferring (**Fig. 2E**) and mouth-preferring neurons (**Fig. 2F**). Given this different pattern of modulation, we next tested whether the electrophysiological properties of eye-preferring and mouth-preferring neurons differed, which might indicate different cell types. However, we found no statistically significant differences in mean firing rate (**Fig. 3A**; eye-preferring: 3.00±3.05 Hz (mean±SD across neurons), mouth-preferring: 3.00±3.58 Hz; two-tailed two-sample *t*-test: *t*(72) = 0.008, P = 0.99) nor in the variability of spike times (see **Methods**), as quantified by the burst index (**Fig. 3B**; eye-preferring: 0.09±0.08, mouth-preferring: 0.10±0.09; *t*(72) = 0.79, P = 0.43) and the modified coefficient-of-variation (CV_2_) (**Fig. 3C, D**; eye-preferring: 1.02±0.114, mouth-preferring: 01.01±0.14; *t*(72) = 0.043, P = 0.97). Moreover, waveforms of eye-preferring and mouth-preferring neurons did not differ significantly (**Fig. 3E, F**) and the trough-to-peak times were statistically indistinguishable (**Fig. 3G, H**; eye-preferring: 0.81±0.22 ms, mouth-preferring: 0.80±0.20 ms; *t*(72) = 0.13, P = 0.90; KS-test: P = 0.97; proportion of neurons with trough-to-peak times > 0.5 ms: eye-preferring: 50/55, mouth-preferring: 17/19; χ^2^-test: P = 0.85). Lastly, neither eye-preferring (**Fig. 3I**; *r*(55) = 0.26, P = 0.050) nor mouth-preferring neurons (**Fig. 3J**; *r*(19) = 0.38, P = 0.11) showed a significant correlation between mean firing rate and waveform as quantified by trough-to-peak time. Together, the basic electrophysiological signatures suggest that eye-preferring and mouth-preferring neurons were not of different neuronal types.

**Fig. 3.**
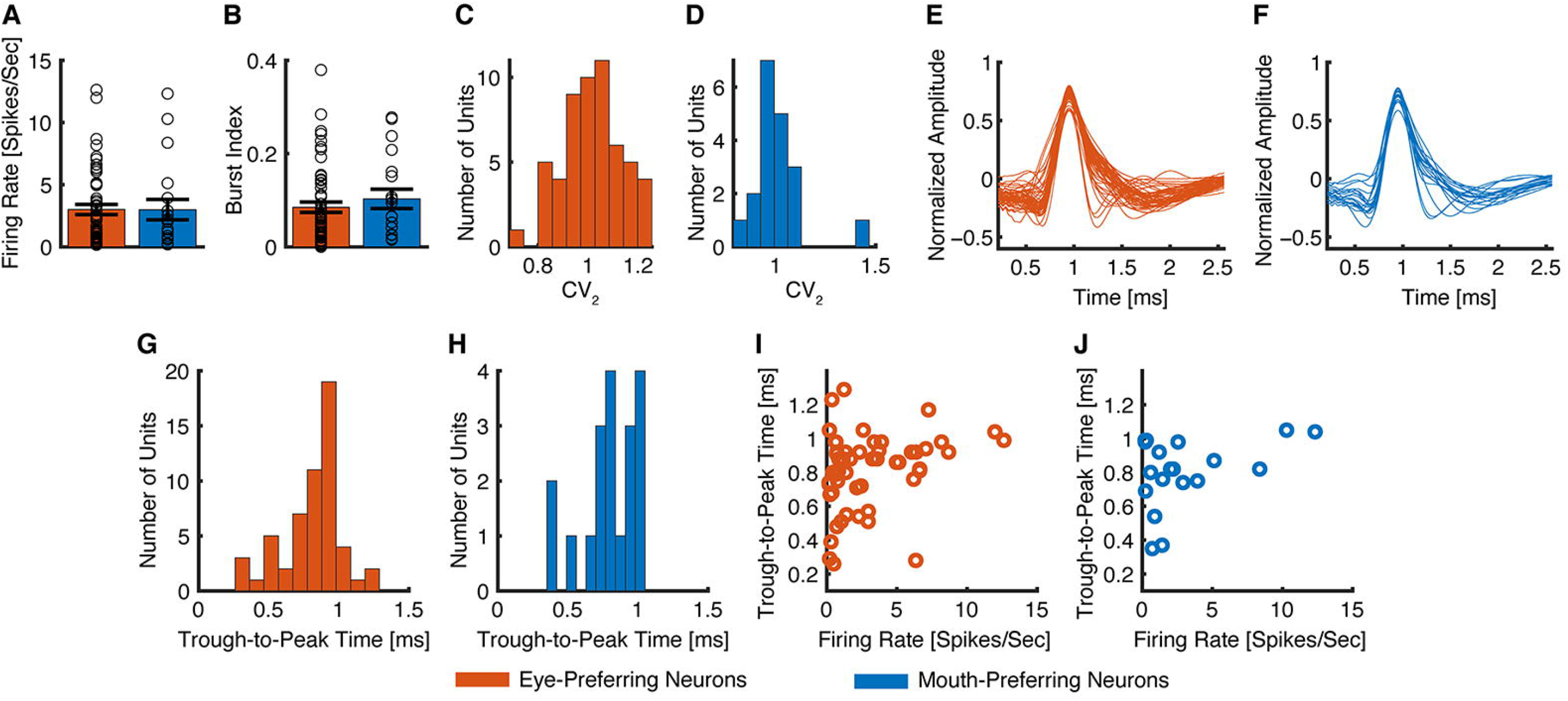
Comparison of cell type between eye-preferring neurons and mouth-preferring neurons. **(A)** Mean firing rate. Error bar denotes ±SEM across neurons and circles show individual values. Red: eye-preferring neurons. Blue: mouth-preferring neurons. **(B)** Burst index (BI). **(C, E, G, I)** Eye-preferring neurons (n = 55). **(D, F, H, J)** Mouth-preferring neurons (n = 19). **(C, D)** Distribution of the modified coefficient-of-variation (CV_2_). **(E, F)** Mean action potential waveforms. **(G, H)** Distribution of trough-to-peak times. **(I, J)** Correlation between mean firing rate and trough-to-peak time. Neither eye-preferring neurons nor mouth-preferring neurons showed a significant correlation.

### Neurons that encoded saccades to the eyes and mouth

We next investigated whether neurons encoded saccades to the eyes or mouth. We aligned neuronal responses at saccade onset and used the mean firing rate in a time window from 200 ms before to 200 ms after saccade onset to calculate statistics. We identified 29 neurons (7.95%; binomial P = 0.006) that had a response differing significantly between saccades to the eyes vs. saccades to the mouth (two-tailed *t*-test, P < 0.05). Among these neurons, 22 (75.9%) had a greater response for saccades to the eyes (see **Fig. 4A, B** for examples and **Fig. 4E** for group results) and 7 (24.1%) had a greater response for saccades to the mouth (see **Fig. 4C, D** for examples and **Fig. 4F** for group results). Similar to fixations, we quantified the response during individual saccades using a saccade-selectivity index (SSI; see **Methods**). As expected, the SSI for selective neurons was significantly larger during saccades to the eyes compared to saccades to the mouth (two-tailed two-sample KS-test, KS = 0.16, P = 8.88×10^−40^; **Fig. 4G**).

**Fig. 4.**
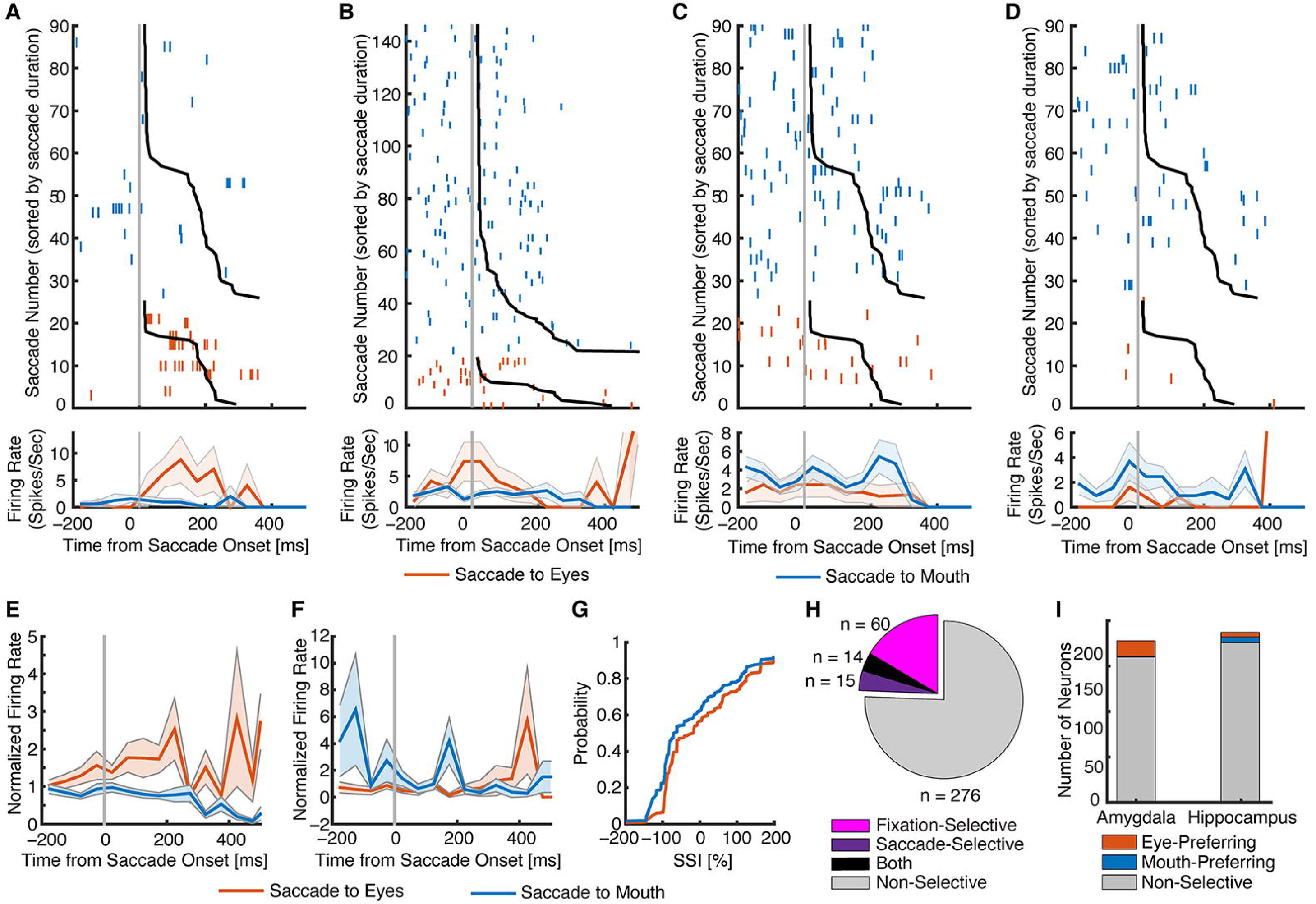
Neurons that encode saccades to eyes vs. mouth. **(A-D)** Saccade-aligned examples. **(A, B)** Neurons that had a greater firing rate when saccading to eyes compared to mouth (selection by two-tailed *t*-test in a time window of −200 ms to 200 ms relative to saccade onset: both Ps < 0.01^=^). **(C, D)** Neurons that had a greater firing rate when saccading to mouth compared to eyes (both Ps < 0.05). Saccades are sorted by saccade duration (black line shows start of the next fixation). t = 0 is saccade onset. Asterisk indicates a significant difference between saccades to eyes / mouth vs. other facial parts in that bin (P < 0.05, two-tailed *t*-test, after Bonferroni correction; bin size = 50 ms). **(E-I)** Population summary. **(E)** Average normalized firing rate of neurons that were selective to saccades to eyes (n = 22). **(F)** Average normalized firing rate of neurons that were selective to saccades to mouth (n = 7). Shaded area denotes ±SEM across neurons. Asterisk indicates a significant difference between the conditions in that bin (P < 0.05, two-tailed *t*-test, after Bonferroni correction). **(G)** Single-saccade analysis using the SSI (**Methods**). Shown is the cumulative distribution of the single-saccade response. **(H)** Overlap between neurons showing fixation selectivity (magenta) and neurons showing saccade selectivity (purple). Black: both. Gray: neither. **(I)** The number of neurons in the amygdala and hippocampus. Stacked bar shows neurons that were selective to saccades to eyes (red), neurons that were selective to saccades to mouth (blue), and non-selective neurons (gray).

Furthermore, we found that 14/29 neurons (48.28%) that differentiated saccades to the eyes vs. the mouth were also feature-selective neurons (i.e., differentiated fixations onto the eyes vs. the mouth), showing a significantly higher percentage than the overall population (20.27%; χ^2^-test: P = 4.92×10^−4^; **Fig. 4H**). This suggests that saccade selectivity was related to subsequent fixation selectivity (similar results were derived when we established fixation selectivity without including the preceding saccade time interval: P = 6.73×10^−4^). Lastly, we derived similar results when we restricted our analysis within saccades between the eyes and mouth (i.e., excluding saccades initiated elsewhere), and we also derived similar results when we compared saccades to the eyes or mouth separately to all other saccades.

Together, we found a population of MTL neurons that encode saccade targets to the eyes and mouth. These neurons may elicit further responses of fixations onto the eyes and mouth.

### Population decoding

How representative of the entire population of recorded MTL neurons are the subsets of neurons described so far? In particular, although non-selective neurons could not distinguish between fixations on the eyes vs. the mouth or between saccades to the eyes vs. the mouth individually, could they still do so as a population? To answer these questions, we next employed population decoding. As expected, decoding from all recorded neurons together and decoding from feature-selective neurons revealed a strong ability to differentiate between fixations on the eyes vs. the mouth (**Fig. 5A**[magenta]; for all time bins). However, this ability was retained in non-feature-selective neurons as well (**Fig. 5A**[gray]; primarily for later time bins), suggesting that non-feature-selective neurons still carried information about differentiating facial features. Similarly, the whole population of neurons and those selective to saccade targets showed a strong ability to differentiate between saccades to the eyes vs. the mouth, but the non-selective neurons also partially retained this ability (**Fig. 5B**; primarily for early time bins).

**Fig. 5.**
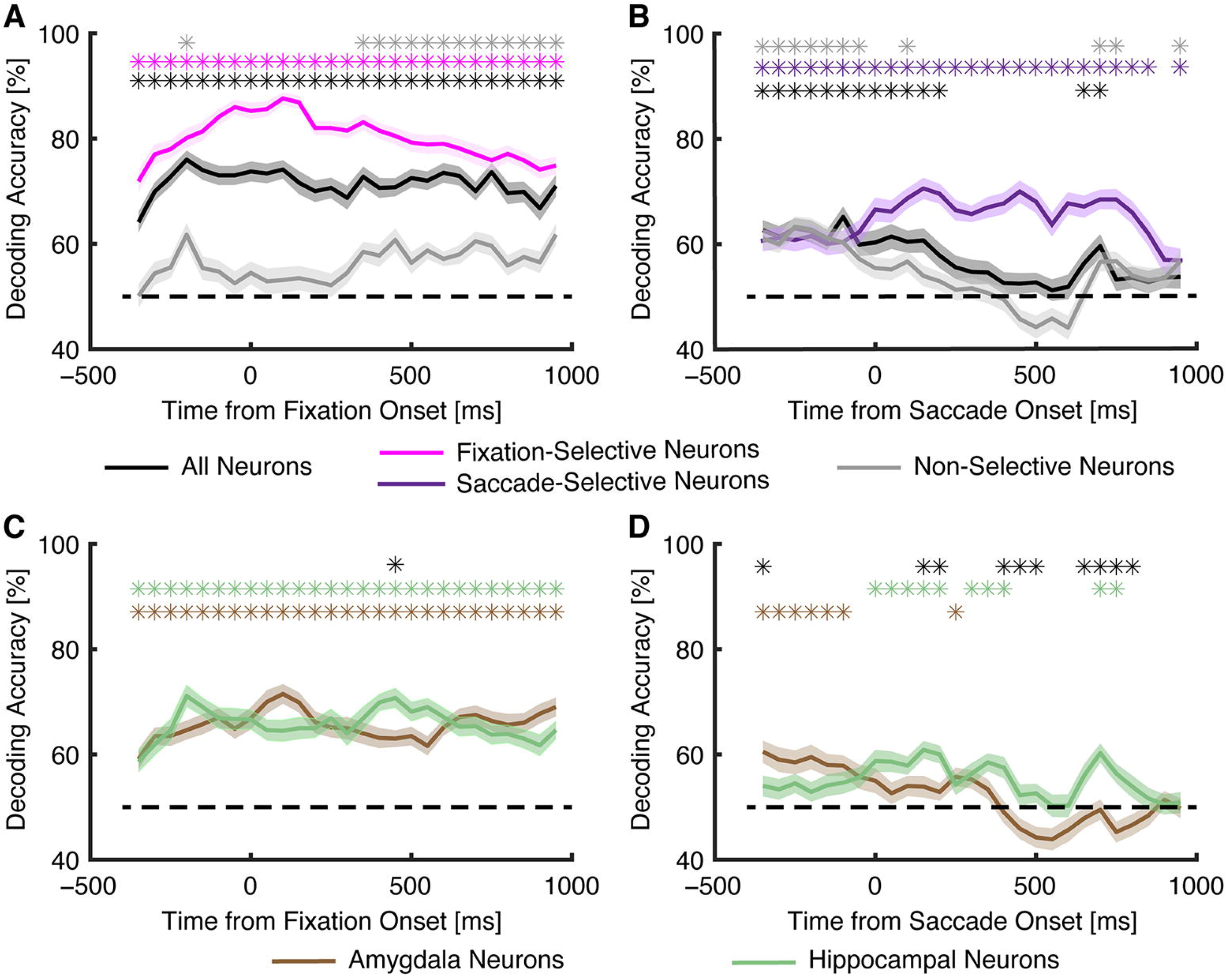
Neuronal population decoding. **(A, C)** Decoding of fixation on the eyes vs. the mouth. Bin size is 500 ms and step size is 50 ms. The first bin is from −600 ms to −100 ms (bin center:−350 ms) relative to fixation onset, and the last bin is from 700 ms to 1200 ms (bin center: 950 ms) after fixation onset. **(B, D)** Decoding of saccade to eyes vs. mouth. Bin size is 500 ms and step size is 50 ms. The first bin is from −600 ms to −100 ms (bin center: −350 ms) relative to fixation onset, and the last bin is from 700 ms to 1200 ms (bin center: 950 ms) after fixation onset. **(A, B)** Decoding with all neurons (black), fixation-selective neurons (magenta), saccade-selective neurons (purple), and non-selective neurons (gray). Shaded area denotes ±SEM across bootstraps. The horizontal dashed black line indicates the chance level (50%). The top asterisks illustrate the time points with a significant above-chance decoding performance (paired *t*-test against chance level, P < 0.05, corrected by FDR for Q < 0.05). **(C, D)** Decoding with all amygdala neurons (brown) and all hippocampal neurons (green). The top brown and green asterisks illustrate the time points with a significant above-chance decoding performance (paired *t*-test against chance level, P < 0.05, corrected by FDR for Q < 0.05). The top black asterisks illustrate the time points with a significant difference between amygdala and hippocampal neurons (two-tailed two-sample *t*-test, P < 0.05, corrected by FDR for Q < 0.05).

### The amygdala and hippocampus played a different role in encoding facial features

Do the amygdala and hippocampus contribute equally to encoding facial features? Notably, we found that the amygdala (47/178 neurons; 26.4%) had a greater percentage of feature-selective neurons compared with the hippocampus (27/187 neurons; 14.4%; χ^2^-test: P = 0.0045; **Fig. 2J**; **Table S1**). Furthermore, we found that among feature-selective neurons, the amygdala had a much greater percentage of eye-preferring neurons (42/47 neurons; 89.4%) than the hippocampus (13/27 neurons; 48.2%; χ^2^-test: P = 9.35×10^−5^; **Fig. 2J**; **Table S1**; and thus the amygdala had a lower percentage of mouth-preferring neurons). Therefore, the amygdala might play a different role in encoding facial features compared with the hippocampus, based on the greater percentage of feature-selective (in particular eye-preferring) neurons in the amygdala. However, feature-selective neurons from the amygdala vs. hippocampus were not of different cell types (**Fig. S2**).

Although the amygdala (18/178 neurons; 10.11%) did not have a significantly greater percentage of neurons that differentiated saccades to the eyes vs. the mouth than the hippocampus (11/187 neurons; 5.88%; χ^2^-test: P = 0.14; **Fig. 4I**), among these selective neurons, the amygdala (17/18 neurons; 94.4%) had a greater percentage of neurons that had a higher response to saccades to the eyes than those in the hippocampus (5/11 neurons; 45.5%; χ^2^-test: P = 0.0028; **Fig. 4I**).

Furthermore, both amygdala neurons and hippocampal neurons showed an above-chance decoding performance for fixation selectivity (**Fig. 5C**; for all time bins) and the decoding performance was similar between the amygdala and hippocampus (**Fig. 5C**). This confirms that both amygdala and hippocampal neurons contained information about fixations onto facial features. Interestingly, amygdala and hippocampal neurons had different decoding temporal dynamics when they encoded saccades (**Fig. 5D**): amygdala neurons tended to encode saccades earlier before saccade onset whereas the hippocampus tended to encode saccades later after saccade onset. This difference may indicate a hierarchal processing of saccades in the human medial temporal lobe.

### Feature-selective neurons encoded the saliency of facial features for face identification

We next analyzed the functional consequence of feature-selective neurons. Using the VGG-16 DNN trained for face recognition (Omkar M. Parkhi 2015), we constructed saliency maps of faces that represented critical features leading to face identification (i.e., the discriminative features of each face identity; **Fig. 6A**). We then correlated the saliency value under each fixation with the corresponding neuronal firing rate for each neuron (**Fig. 6B**). We found that feature-selective neurons showed a significantly above-chance (0) correlation for fixations within the preferred ROIs (i.e., within eyes and mouth; Pearson’s *r* = 0.04±0.07; mean±SD across neurons; two-tailed paired *t*-test against 0: *t*(73) = 4.28, P = 5.52×10^−5^) but not for fixations outside the preferred ROIs (*r* = 0.015±0.08; *t*(73) = 1.51, P = 0.13; **Fig. 6C**). Note that saliency was analyzed separately within or outside the eyes/mouth ROIs because these ROIs tended to have a higher saliency value (e.g., **Fig. 6A**; in other words, eyes/mouth contained more information about face recognition) and feature-selective neurons could discriminate eyes/mouth from other facial parts (**Fig. 2G**). Furthermore, feature-selective neurons showed a greater correlation with saliency values than non-feature-selective neurons for fixations within the eyes/mouth ROIs (**Fig. 6C**; *t*(363) = 2.25, P = 0.025) and but not for fixations outside the eyes/mouth ROIs (*t*(363) = 1.18, P = 0.24). Together, our results suggest that feature selectivity of eyes vs. mouth is related to face recognition, a possible functional consequence of feature-selective neurons.

**Fig. 6.**
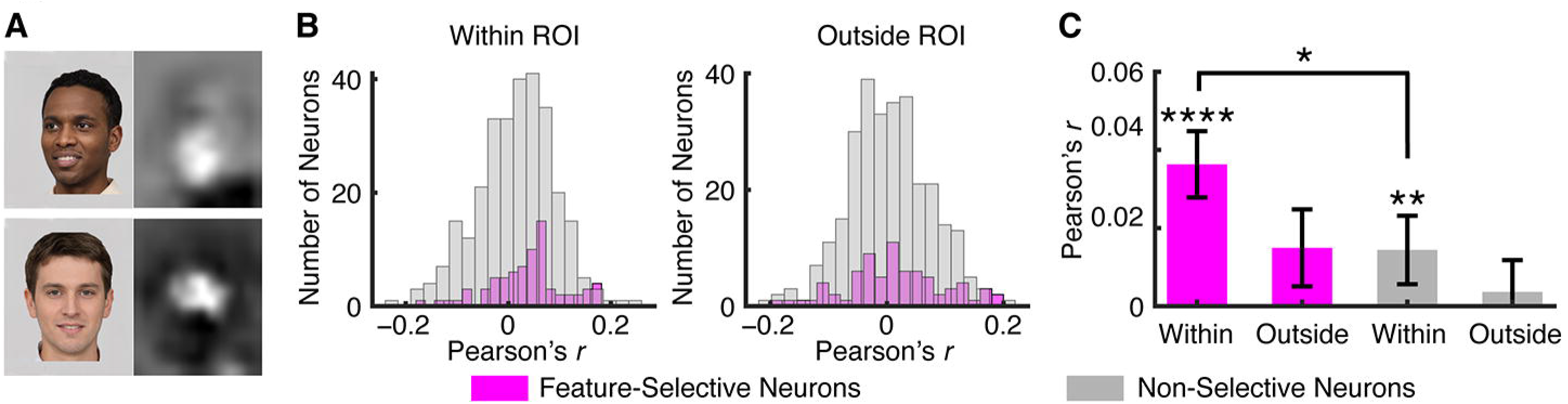
Feature-selective neurons encoded the saliency of facial features for face identification. **(A**) Saliency maps (right) for two sample face stimuli (left). **(B)** Distribution of correlation coefficients (Pearson’s *r*) across neurons. Correlation was calculated between the firing rate and saliency value across fixations for each neuron. The ROI combined both the eyes and the mouth. **(C)** Neuronal population average within and outside of the ROI. Error bars denote ±SEM across neurons. Asterisks indicate a significant difference above 0 (two-tailed paired *t*-test) or between feature-selective vs. non-selective neurons (two-tailed two-sample *t*-test). *: P < 0.05, **: P < 0.01, and ****: P < 0.0001.

### Feature-selective neurons correlated with perceived social trait judgment

Lastly, we analyzed whether feature-selective neurons were related to implicit judgment of social traits. Because patients passively viewed the faces without providing explicit judgment of faces during neural recordings, social traits might be implicitly processed in the brain. We subsequently acquired explicit ratings on a subset of faces (2 to 5 faces for each identity) through online questionnaires after patients were discharged. We acquired ratings for the following 8 social traits (Lin et al 2019): *warm, critical, competent, practical, feminine, strong, youthful*, and *charismatic*, and we used the average rating across patients for each face to calculate correlations. We found that the firing rate for fixations on the eyes correlated with the social trait of *warmth*, for both feature-selective neurons (Pearson’s *r* = −0.03±0.06; mean±SD across neurons; two-tailed paired *t*-test against 0: *t*(73) = 4.05, P = 0.001; **Fig. 7A**) and non-feature-selective neurons (*r* = −0.011±0.06; *t*(288) = 3.23, P = 0.0014), but feature-selective neurons showed a stronger correlation (*t*(361) = 2.43, P = 0.02). Similarly, the firing rate for fixations on the eyes correlated with the social trait of *practical* (feature-selective neurons: *r* = −0.028±0.066; *t*(73) = 3.60, P = 5.84×10^−4^; non-feature-selective neurons: *r* = 0.003±0.064; *t*(288) = 0.77, P = 0.44; **Fig. 7A**), and feature-selective neurons showed a stronger correlation (*t*(361) = 3.64, P = 3.13×10^−4^). Furthermore, we found that the firing rate for fixations on the mouth correlated with the social trait of *feminine* (feature-selective neurons: *r* = −0.03±0.07; *t*(64) = 2.99, P = 0.004; non-feature-selective neurons: *r* = 0.006±0.06; *t*(273) = 1.45, P = 0.15; **Fig. 7B**) and *strong* (feature-selective neurons: *r* = 0.026±0.09; *t*(64) = 2.35, P = 0.02; non-feature-selective neurons: *r* = 0.006±0.07; *t*(273) = 1.45, P = 0.15; **Fig. 7B**), and for both traits (*feminine* and *strong*), feature-selective neurons showed a stronger correlation (feminine: *t*(337) = 2.26, P = 0.02; strong: *t*(337) = 1.97, P = 0.0498).

**Fig. 7.**
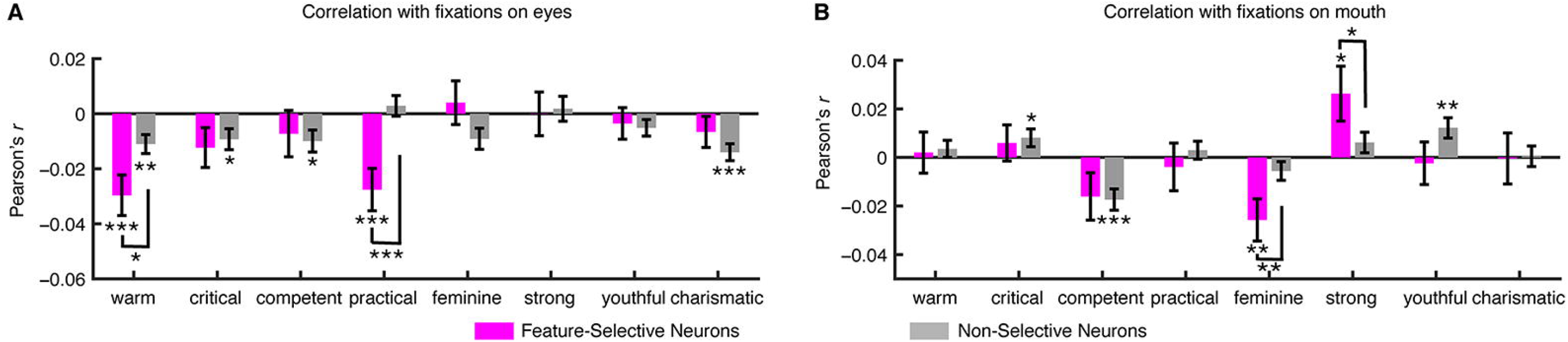
Feature-selective neurons encoded social traits. **(A)** Correlation between the firing rate for fixations on the eyes and perceived social traits. **(B)** Correlation between the firing rate for fixations on the mouth and perceived social traits. Error bars denote ±SEM across neurons. Asterisks indicate a significant difference above 0 (two-tailed paired *t*-test) or between feature-selective vs. non-selective neurons (two-tailed two-sample *t*-test). *: P < 0.05, **: P < 0.01, and ***: P < 0.001.

Together, we found that the neuronal response to facial features was related to perception of social traits and feature-selective neurons encoded these social traits more strongly than non-feature-selective neurons.

## Discussion

In this study, we recorded neural activity from amygdala and hippocampal neurons in neurosurgical patients using implanted depth electrodes while they viewed static images of famous faces. We identified the neural mechanisms underlying human visual scanning patterns of faces by revealing a class of neurons sensitive to different facial features (the eyes and mouth). We found that the amygdala had a greater percentage of feature-selective neurons that were eye-preferring compared with the hippocampus, although feature-selective neurons from the amygdala vs. hippocampus and eye-preferring vs. mouth-preferring neurons were not of different neuronal types. We further revealed neurons that were sensitive to saccade targets and we confirmed these results using population decoding. Lastly, we revealed functional roles of feature-selective neurons. Feature-selective neurons encoded the saliency of facial features so they might contribute to face identification; and feature-selective neurons were correlated with perceived social traits so they might also contribute to face evaluation of social judgments. Together, we not only characterized a new class of neurons that encoded facial features, but also suggest they may play important roles in face recognition and judgment.

Previous neuroimaging studies have shown that the amygdala is sensitive to facial features and saccade directions (Gamer & Büchel 2009), consistent with our present findings. These neuroimaging studies varied face locations on the screen for each trial (either aligning the eyes or mouth to the screen center) in order to control for fixation starting location and fixation sequence. In the present study, our single-neuron recordings had a superior temporal resolution and our simultaneous eye tracking allowed fixation-based analysis (rather than trial-based found in neuroimaging studies). We have further shown that our observed feature selectivity could not be explained by fixation serial order (**Fig. 2I**). Furthermore, compared with previous research, our findings were derived using a large variety of natural face images; and we have replicated our findings using separate tasks and a new set of cartoon faces (manuscript in preparation). In addition, we have extended previous research showing that amygdala neurons are selective to whole vs. piecemeal faces (Rutishauser et al 2011) by further showing that amygdala and hippocampal neurons are selective to fixations and saccades to salient facial features such as the eyes and mouth.

We found feature-selective neurons in both the amygdala and hippocampus but the amygdala had a higher percentage of feature-selective neurons and in particular eye-preferring neurons than the hippocampus, which highlighted a different role between the amygdala and hippocampus in encoding facial features. Such difference in function between the amygdala and hippocampus is consistent with our previous finding showing that only the amygdala but not the hippocampus encode perceived facial emotions (Wang et al 2014). However, the amygdala and hippocampus play a similar role in encoding face identities (Cao et al 2020), visual selectivity (Kreiman et al 2000, Wang et al 2018), memory (Rutishauser et al 2010), and visual attention (Wang et al 2018). Furthermore, although some faces in the CelebA stimuli have facial expressions (because the celebrities posed when taking the photos) and eye movement can be biased by facial expressions (Scheller et al 2012), we observed similar results in an unpublished study using emotionally neutral FaceGen model faces. Therefore, our results could not be simply explained by the neurons’ response to facial emotions (Wang et al 2014, Wang et al 2017).

## Methods

### Participants

There were 16 sessions from 5 patients in total (**Table S1**). All sessions had simultaneous eye tracking. We excluded 3 sessions that had fewer than 10 fixations onto each facial region of interest (ROI), resulting in a total of 13 sessions for further analysis. All participants provided written informed consent using procedures approved by the Internal Review Board of the West Virginia University (WVU).

### Stimuli

We used faces of celebrities from the CelebA dataset (Liu et al 2015). We selected 50 identities with 10 images for each identity, totaling 500 face images. The identities were selected to include both genders and multiple races. We used the same stimuli for all patients. Patients were asked to indicate whether they were familiar with each identity in a follow-up survey.

Patients were also asked to provide judgments of social traits on a 1 to 7 scale through an online questionnaire after they were discharged. The social traits include *warm, critical, competent, practical, feminine, strong, youthful*, and *charismatic*. Three patients completed the questionnaire and depending on the availability of the patients, patients provided ratings for 2 to 5 faces per identity per social trait (the rated faces were all from the original stimuli). We observed high inter-subject consistency so we used the average rating for each face to correlate perceptions of social judgment with neuronal firing rate. We also included neuronal data from the two patients who did not provide ratings.

### Experimental procedure

We used a simple 1-back task. In each trial, a single face was presented at the center of the screen for a fixed duration of 1 second, with uniformly jittered inter-stimulus-interval (ISI) of 0.5-0.75 seconds (**Fig. 1A**). Each image subtended a visual angle of approximately 10°. Patients pressed a button if the present face image was *identical* to the immediately previous image. 9% of trials were one-back repetitions. Each face was shown once unless repeated in one-back trials; and we excluded responses from one-back trials to have an equal number of responses for each face. This task kept patients attending to the faces, but avoided potential biases from focusing on a particular facial feature (e.g., compared to asking patients to judge a particular facial feature). The order of faces was randomized for each patient. This task procedure has been shown to be effective to study face representation in humans (Grossman et al 2019).

Stimuli were presented using MATLAB with the Psychtoolbox 3 (Brainard 1997) (http://psychtoolbox.org) (screen resolution: 1600 × 1280).

### Eye tracking

Patients were recorded with a remote non-invasive infrared Eyelink 1000 system (SR Research, Canada). One of the eyes was tracked at 500 Hz. The eye tracker was calibrated with the built-in 9-point grid method at the beginning of each block. Fixation extraction was carried out using software supplied with the Eyelink eye tracking system. Saccade detection required a deflection of greater than 0.1°, with a minimum velocity of 30°/s and a minimum acceleration of 8000°/s^2^, maintained for at least 4 ms. Fixations were defined as the complement of a saccade, i.e. periods without saccades. Analysis of the eye movement record was carried out off-line after completion of the experiments.

### Electrophysiology

We recorded from implanted depth electrodes in the amygdala and hippocampus from patients with pharmacologically intractable epilepsy. Target locations in the amygdala and hippocampus were verified using post-implantation CT. At each site, we recorded from eight 40 μm microwires inserted into a clinical electrode as described previously (Rutishauser et al 2006a, Rutishauser et al 2010). Efforts were always made to avoid passing the electrode through a sulcus, and its attendant sulcal blood vessels, and thus the location varied but was always well within the body of the targeted area. Microwires projected medially out at the end of the depth electrode and examination of the microwires after removal suggests a spread of about 20-30 degrees. The amygdala electrodes were likely sampling neurons in the mid-medial part of the amygdala and the most likely microwire location is the basomedial nucleus or possibly the deepest part of the basolateral nucleus. Bipolar wide-band recordings (0.1-9 kHz), using one of the eight microwires as the reference, were sampled at 32 kHz and stored continuously for off-line analysis with a Neuralynx system. The raw signal was filtered with zero-phase lag 300-3 kHz bandpass filter and spikes were sorted using a semi-automatic template matching algorithm as described previously (Rutishauser et al 2006b). Units were carefully isolated and recording and spike sorting quality were assessed quantitatively (**Fig. S1**).

### Spikes

Only units with an average firing rate of at least 0.15 Hz (entire task) were considered (Cao et al 2020). Fixations were aligned to fixation onset and saccades were aligned to saccade onset. Average firing rates Peri-Stimulus-Time Histogram (PSTH) were computed by counting spikes across all fixations or across all saccades in consecutive 50 ms bins. Pairwise comparisons were made using a two-tailed *t*-test at P < 0.05 and Bonferroni-corrected for multiple comparisons in time bins in the group PSTH. Asterisks in the figures indicate a significant difference after Bonferroni correction.

### Data analysis: response index for single fixation or saccade

For each neuron we quantified whether its response differed between fixation on the eyes and fixation on the mouth using a single-fixation selectivity index, FSI **(Eq. 1)**. The FSI facilitates group analysis and comparisons between different types of cells (i.e., eye- and mouth-preferring cells in this study), as motivated by previous studies (Wang et al 2018, Wang et al 2014). The FSI quantifies the response during fixation *i* relative to the mean response to fixations on the mouth and baseline (the interval right before face onset). The mean response and baseline were calculated individually for each neuron.

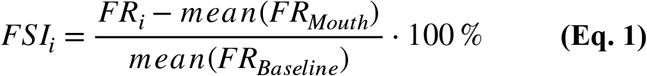

For each fixation *i*, *FSI*_*i*_ is the baseline normalized mean firing rate (FR) during an interval from 200 ms before fixation onset to 200 ms after fixation offset (the same time interval as cell selection). Different time intervals were tested as well, to ensure that results were qualitatively the same and not biased by particular spike bins.

If a neuron distinguishes fixations on the eyes from fixations on the mouth, the average value of *FSI*_*i*_ of all fixations will be significantly different from 0. Since eye-preferring neurons have more spikes in fixations on the eyes and mouth-preferring neurons have more spikes in fixations on the mouth, on average *FSI*_*i*_ is positive for eye-preferring neurons and negative for mouth-preferring neurons. To get an aggregate measure of activity that pools across neurons, *FSI*_*i*_ was multiplied by −1 if the *neuron* is classified as a mouth-preferring neuron **(Eq. 2)**. This makes *FSI*_*i*_ on average positive for both types of feature-selective neurons. Notice that the factor −1 depends only on the *neuron* type but not fixation type. Thus, negative *FSI*_*i*_ values are still possible.

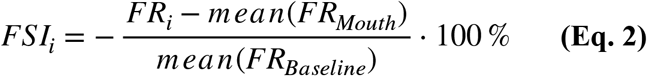

After calculating *FSI*_*i*_ for every fixation, we subsequently averaged all *FSI*_*i*_ of fixations that belong to the same category. By definition, the average value of *FSI*_*i*_ for fixation on the mouth will be equal to zero because the definition of *FSI*_*i*_ is relative to the response to fixation on the mouth (see **Eq. 2**). The mean baseline firing rate was calculated across all trials. The same *FR*_*Mouth*_ was subtracted for both types of fixations.

The cumulative distribution function (CDF) was constructed by calculating for each possible value *x* of the FSI how many examples are smaller than x. That is, F(*x*) = P(X ≤ x), where X is a vector of all FSI values. The CDF of fixations on the eyes and mouth were compared using two-tailed two-sample Kolmogorov-Smirnov (KS) tests.

Similarly, we defined single-saccade selectivity index (SSI) during an interval from 200 ms before to 200 ms after saccade onset (the same time interval as cell selection) as we quantified saccades to the eyes or mouth.

### Data analysis: single-neuron ROC analysis

Neuronal Receiver Operating Characteristics (ROCs) were constructed based on the spike counts in a time window of 200 ms before fixation onset to 200 ms after fixation offset for fixation-wise analysis. We varied the detection threshold between the minimal and maximal spike count observed, linearly spaced in 20 steps. The Area Under the Curve (AUC) of the ROC was calculated by integrating the area under the ROC curve (trapezoid rule). The AUC value is an unbiased estimate for the sensitivity of an ideal observer that counts spikes and makes a binary decision based on whether the number of spikes is above or below a threshold. We defined the category with a higher overall firing rate as ‘true positive’ and the category with a lower overall firing rate as ‘false positive’. Therefore, the AUC value was always above 0.5 by definition.

### Data analysis: neuronal population decoding

We pooled all recorded neurons into a large pseudo-population (see (Rutishauser et al 2015, Wang et al 2019)). Firing rates were *z*-scored individually for each neuron to give equal weight to each unit regardless of firing rate. We used a maximal correlation coefficient classifier (MCC) as implemented in the MATLAB neural decoding toolbox (NDT) (Meyers 2013). The MCC estimates a mean template for each class *i* and assigns the class for test fixation. We used 8-fold cross-validation, i.e., all fixations were randomly partitioned into 8 equal sized subsamples, of which 7 subsamples were used as the training data and the remaining single subsample was retained as the validation data for assessing the accuracy of the model, and this process was repeated 8 times, with each of the 8 subsamples used exactly once as the validation data. We then repeated the cross-validation procedure 50 times for different random train/test splits. Statistical significance of the decoding performance for each group of neurons against chance was estimated by calculating the percentage of bootstrap runs (50 in total) that had an accuracy below chance (i.e., 50% when decoding the type of ROI). Statistical significance for comparing between groups of neurons was estimated by calculating the percentage of bootstrap runs (50 in total) that one group of neurons had a greater accuracy than the other. For both tests, we used false discovery rate (FDR) (Benjamini & Hochberg 1995) to correct for multiple comparisons across time points. Spikes were counted in bins of 500 ms size and advanced by a step size of 50 ms. The first bin started −600 ms relative to fixation onset (bin center was thus 350 ms before fixation onset), and we tested 27 consecutive bins (the last bin was thus from 700 ms to 1200 ms after fixation onset). For each bin, a different classifier was trained/tested.

For decoding of saccades, we also used 8-fold cross-validation and repeated the process 50 times with different subsets of saccades, resulting in a total of 400 tests to estimate the test performance. Spikes were counted in bins of 500 ms size and advanced by a step size of 50 ms. The first bin started −600 ms relative to saccade onset (bin center was thus 350 ms before saccade onset), and we tested 27 consecutive bins (the last bin was thus from 700 ms to 1200 ms after saccade onset). For each bin, a different classifier was trained/tested.

### Data analysis: comparison of cell types

We quantified basic electrophysiological parameters following previous studies (Rutishauser et al 2013, Viskontas et al 2007). To compare the variability of spike times, we computed the inter-spike interval (ISI) distribution of each cell by considering all spikes fired during the experiment and quantified it using two metrics: the burst index (BI) and the modified coefficient-of-variation (CV_2_). The BI was defined as the proportion of ISIs less than 10 ms (Wyler et al 1975). The CV_2_ (**Eq. 3**) is a function of the difference between two adjacent ISIs and is a standard measure to quantify spike-train variability that is robust to underlying rate changes (Holt et al 1996). In contrast, the coefficient-of-variation measure CV is only valid for stationary processes (i.e., fixed mean firing rate) and is thus not applicable for this analysis.

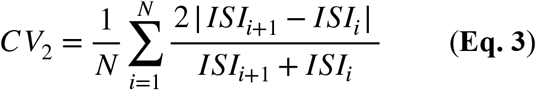

We compared the waveform of different neurons based on the trough-to-peak time of the mean waveform (Mitchell et al 2007). The mean waveform is the average of all spikes assigned to the cluster. The polarity of the mean waveforms was inverted if necessary such that the trough always occurs before the peak. We also evaluated whether there was a correlation between the trough-to-peak time and the mean firing rate of a unit. For this, the mean firing rate was defined as the mean rate over the entire duration of all valid trials.

## Acknowledgements

We thank all patients for their participation, staff from WVU Ruby Memorial Hospital for support with patient testing, and Paula Webster for valuable comments. This research was supported by an NSF CAREER Award (1945230), ORAU Ralph E. Powe Junior Faculty Enhancement Award, West Virginia University (WVU), WVU PSCoR Program, and the Dana Foundation (to S.W.), and an NSF Grant (OAC-1839909) and the WV Higher Education Policy Commission Grant (HEPC.dsr.18.5) (to X.L.). The funders had no role in study design, data collection and analysis, decision to publish, or preparation of the manuscript.

## Author Contributions

R.C., X.L., and S.W. designed research. R.C. and S.W. performed experiments. N.J.B. performed surgery. R.C. and S.W. analyzed data. R.C, X.L., and S.W. wrote the paper. All authors discussed the results and contributed toward the manuscript.

---

**Table S1.** List of patients. Each row of neurons represents a separate recording session using. Each session was recorded on a separate day. Total: all neurons recorded from an area. Left: neurons that were recorded from the left hemisphere and had a firing rate greater than 0.15 Hz. Right: neurons that were recorded from the right hemisphere and had a firing rate greater than 0.15 Hz. These neurons were included for further analysis. Eye: eye-preferring feature-selective neurons. Mouth: mouth-preferring feature-selective neurons. Patient P8 was not included in the study because the patient only had surface electrodes.

| ID | Age | Sex | Epilepsy diagnosis | Number of Amygdala Neurons |  |  |  |  | Number of Hippocampal Neurons |  |  |  |  |
| --- | --- | --- | --- | --- | --- | --- | --- | --- | --- | --- | --- | --- | --- |
|  |  |  |  | Total | Left | Right | Eye | Mouth | Total | Left | Right | Eye | Mouth |
| P6 | 33 | F | Left posterior temporal/<br>parietal | 9 | 9 | 0 | 0 | 0 | 31 | 31 | 0 | 1 | 2 |
|  |  |  |  | 10 | 10 | 0 | 0 | 1 | 29 | 29 | 0 | 4 | 4 |
| P7 | 28 | F | Right mesial temporal | 6 | 6 | 0 | 0 | 0 | 21 | 19 | 2 | 0 | 2 |
|  |  |  |  | 6 | 6 | 0 | 1 | 1 | 20 | 19 | 1 | 1 | 0 |
|  |  |  |  | 2 | 2 | 0 | 0 | 0 | 31 | 26 | 5 | 2 | 4 |
| P9 | 42 | M | Left frontal | 28 | 28 | 0 | 4 | 3 | 7 | 7 | 0 | 2 | 1 |
|  |  |  |  | 29 | 29 | 0 | 5 | 0 | 7 | 7 | 0 | 1 | 0 |
|  |  |  |  | 25 | 25 | 0 | 10 | 0 | 6 | 6 | 0 | 0 | 0 |
|  |  |  |  | 23 | 23 | 0 | 15 | 0 | 3 | 3 | 0 | 1 | 0 |
| P10 | 47 | F | Right temporal | 25 | 0 | 25 | 2 | 0 | 7 | 7 | 0 | 1 | 0 |
| P11 | 33 | F | Right temporal | 9 | 0 | 9 | 5 | 0 | 15 | 0 | 15 | 0 | 1 |
|  |  |  |  | 6 | 0 | 6 | 0 | 0 | 10 | 0 | 10 | 0 | 0 |
| Sum |  |  |  | 178 | 138 | 40 | 42 | 5 | 187 | 154 | 33 | 13 | 14 |

## Supplemental Figure Legends

**Fig. S1.** Spike sorting and recording quality assessment. **(A)** Histogram of the number of units identified on each active wire (only wires with at least one unit identified are counted). The average yield per wire with at least one unit was 2.74±1.60 (mean±SD). **(B)** Histogram of mean firing rates. **(C)** Histogram of proportion of inter-spike intervals (ISIs) which are shorter than 3 ms. The large majority of clusters had less than 0.5% of such short ISIs. **(D)** Histogram of the signal-to-noise ratio (SNR) of the mean waveform peak of each unit. **(E)** Histogram of the SNR of the entire waveform of all units. **(F)** Pairwise distance between all possible pairs of units on all wires where more than 1 cluster was isolated. Distances are expressed in units of standard deviation (SD) after normalizing the data such that the distribution of waveforms around their mean is equal to 1. **(G)** Isolation distance of all units for which this metric was defined (n = 365, median = 13.64). **(H)** Eye-preferring and mouth-preferring neurons did not differ significantly in isolation distance (*t*(60) = 1.55, P = 0.13).

**Fig. S2.** Comparison of cell types between amygdala vs. hippocampus feature-selective neurons. **(A)** Mean firing rate. Error bar denotes ±SEM across neurons and circles show individual values. Brown: amygdala feature-selective neurons. Green: hippocampus feature-selective neurons. **(B)** Burst index (BI). **(C, E, G, I)** Amygdala feature-selective neurons (n = 47). **(D, F, H, J)** Hippocampus feature-selective neurons (n = 27). **(C, D)** Distribution of the modified coefficient-of-variation (CV_2_). **(E, F)** Mean action potential waveforms. **(G, H)** Distribution of trough-to-peak times. **(I, J)** Correlation between mean firing rate and trough-to-peak time.

