## Supplementary Materials for "Encoding of facial features by single neurons in the human amygdala and hippocampus"

**Supplemental Figures and Legends**

Figure S1

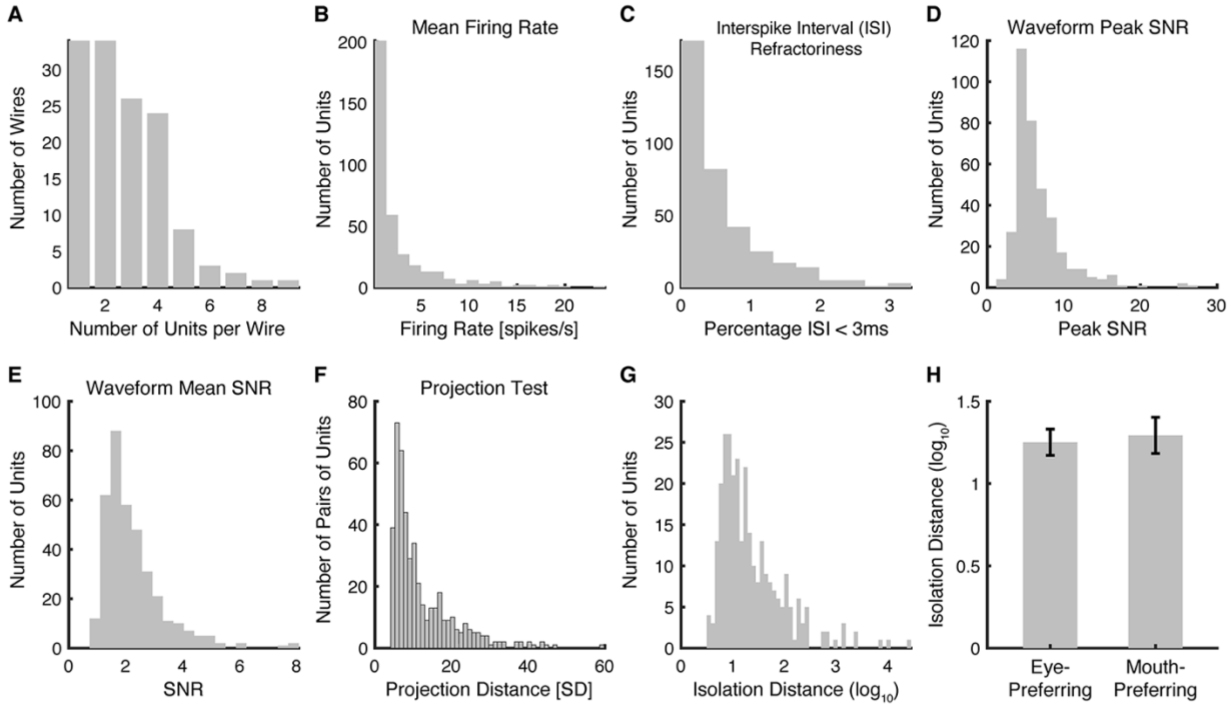


**Fig. S1.** Spike sorting and recording quality assessment. **(A)** Histogram of the number of units identified on each active wire (only wires with at least one unit identified are counted). The average yield per wire with at least one unit was 2.74±1.60 (mean±SD). **(B)** Histogram of mean firing rates. **(C)** Histogram of proportion of inter-spike intervals (ISIs) which are shorter than 3 ms. The large majority of clusters had less than 0.5% of such short ISIs. **(D)** Histogram of the signal-to-noise ratio (SNR) of the mean waveform peak of each unit. **(E)** Histogram of the SNR of the entire waveform of all units. **(F)** Pairwise distance between all possible pairs of units on all wires where more than 1 cluster was isolated. Distances are expressed in units of standard deviation (SD) after normalizing the data such that the distribution of waveforms around their mean is equal to 1. **(G)** Isolation distance of all units for which this metric was defined (n = 365, median = 13.64). **(H)** Eye-preferring and mouth-preferring neurons did not differ significantly in isolation distance ($t(60) = 1.55$, P = 0.13).

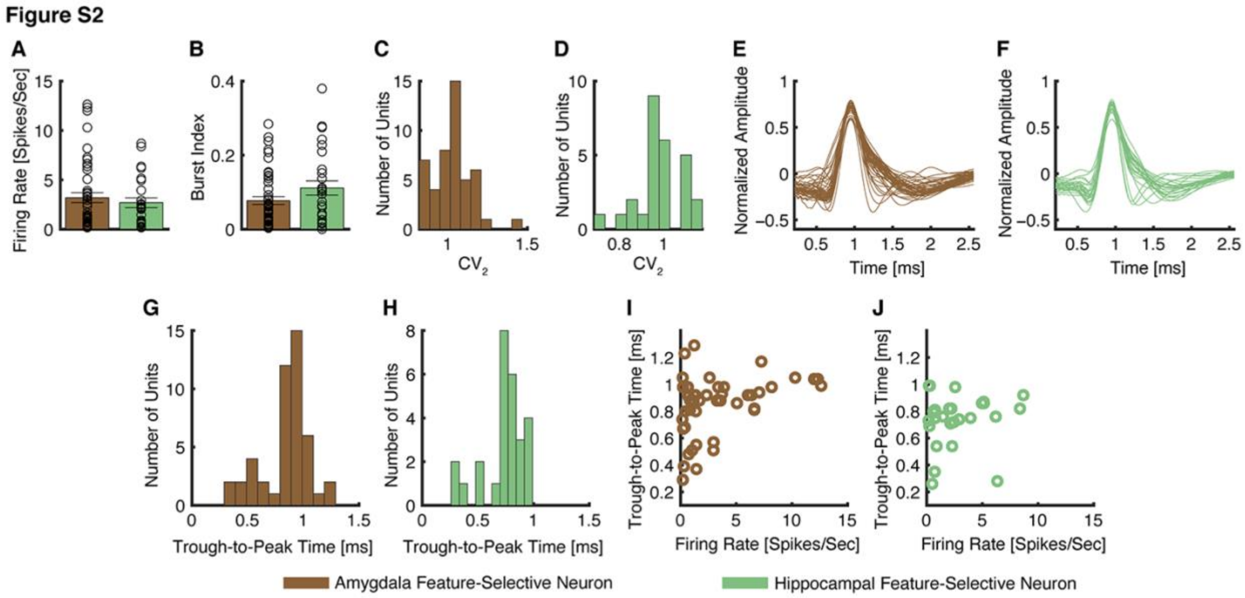


**Fig. S2.** Comparison of cell types between amygdala vs. hippocampus feature-selective neurons. **(A)** Mean firing rate. Error bar denotes ±SEM across neurons and circles show individual values. Brown: amygdala feature-selective neurons. Green: hippocampus feature-selective neurons. **(B)** Burst index (BI). **(C, E, G, I)** Amygdala feature-selective neurons (n = 47). **(D, F, H, J)** Hippocampus feature-selective neurons (n = 27). **(C, D)** Distribution of the modified coefficient-of-variation ($CV_2$). **(E, F)** Mean action potential waveforms. **(G, H)** Distribution of trough-to-peak times. **(I, J)** Correlation between mean firing rate and trough-to-peak time.
